# Darwin’s Overlooked Radiation: genomic evidence points to the early stages of a radiation in the Galápagos prickly pear cactus (*Opuntia,* Cactaceae)

**DOI:** 10.1101/2023.10.03.560736

**Authors:** Felipe Zapata, José Cerca, Dana McCarney, Claudia L. Henriquez, Bashir B. Tiamiyu, John E. McCormack, Kelsey R. Reckling, Jaime A. Chaves, Gonzalo Rivas-Torres

**Affiliations:** Department of Ecology and Evolutionary Biology, University of California, Los Angeles, CA 90095, USA; Center for Tropical Research, Institute of the Environment and Sustainability, University of California, Los Angeles, CA 90095, USA; Center for Ecological and Evolutionary Synthesis (CEES), Department of Biosciences, University of Oslo, Oslo, Norway; School of Medicine, Tufts University, Boston, MA 02155, USA; Department of Plant Biology, University of Ilorin, Nigeria; Moore Laboratory of Zoology, Occidental College, Los Angeles, CA 90041, USA; Department of Biology, San Francisco State University, San Francisco, CA 94132, USA; Colegio de Ciencias Biológicas y Ambientales and Galápagos Institute for the Arts and Sciences, Universidad de San Francisco de Quito USFQ, Quito 170901, Ecuador; Galápagos Science Center UNC—USFQ, San Cristóbal, Galápagos, Ecuador; Department of Geography and Environment, University of North Carolina at Chapel Hill, Chapel Hill, NC 27599, USA

**Author notes:** Correspondence (F.Z.), (J.C.), (G.R.T.). Equal Contribution (Felipe Zapata, José Cerca).

## Abstract

The Galápagos Islands are a prime example of a natural laboratory for the study of evolutionary radiations. While much attention has been devoted to iconic species like Darwin’s finches ^1–4^, the islands offer an equally unique but often overlooked opportunity for plant radiations ^5^. Yet, compared to their animal counterparts, our understanding of the patterns and processes underpinning Galápagos plant radiations remains relatively limited ^6,7^. We present evidence of the early stages of a radiation in prickly-pear cactus (*Opuntia*, Cactaceae), a plant lineage widespread across the archipelago. Phylogenomic and population genomic analyses show that notwithstanding overall low genetic differentiation across populations, there is marked geographic structure that is broadly consistent with current taxonomy and the dynamic paleogeography of the Galápagos. Because such low genetic differentiation stands in stark contrast to the exceptional eco-phenotypic diversity displayed by cacti across islands, it is plausible that phenotypic plasticity precedes genetic divergence and is the source of adaptive evolution, or that introgression between populations facilitates local adaptation. Models of population relationships including admixture indicate that gene flow is common between certain islands, likely facilitated by dispersal via animals known to feed on *Opuntia* flowers, fruits, and seeds across the archipelago. Scans of genetic differentiation between populations reveal candidate loci associated with seed traits and environmental stressors, suggesting that a combination of biotic interactions and abiotic pressures due to the harsh conditions characterizing island life in a volcanic, equatorial archipelago may underlie the diversification of prickly-pear cacti. Considered in concert, these results are relevant to both the mechanisms of plant eco-phenotypic differentiation and the evolutionary history and conservation of the Galápagos biota.

## RESULTS AND DISCUSSION

> ‘‘ I certainly recognise S. America in ornithology, would a botanist? ”
>
> C. R. Darwin (1835, Field Notes, p 30) soon after arriving to the Galápagos and noticing the mockingbirds, which he could associate with birds from mainland South America. He doubted the same could be said about the plants.

The origins, persistence, and diversification of species have been studied extensively in oceanic islands where ecological communities are often less complex and environmental conditions provide variation for evolution to flourish ^1,8–10^. The Galápagos Islands are a prime example of such natural laboratories for studying evolution. Studies on the diversification of Galápagos tortoises ^11,12^, iguanas ^13,14^, land snails ^15^, and Darwin’s finches ^2,16–19^ have provided fundamental biological insight about foundational ideas in evolutionary biology, including speciation, natural selection, and adaptive radiation.

Strikingly, and in spite of a relatively well-known flora, studies about the diversification of Galápagos plant lineages have received considerably less attention, with the exception of a few classic studies on prickly-pear cacti ^20–22^ and recent work on the salt bush ^23^ and the Darwin’s Giant Daisies ^6,7^. This is surprising given the central role that the plants from the Galápagos, particularly the cacti, played in shaping Darwin’s ideas for his theory of evolution ^5^ and in the dietary ecology and diversification of the iconic Darwin’s finches ^2,3,19,24,25^.

Here, we use genomic data to revisit classic work on the prickly-pear cacti (*Opuntia*) across the Galápagos archipelago to investigate the evolution of this charismatic lineage of the Galápagos flora (Fig. 1). Given that these plants furnish an important source of food and water to multiple native animals across the archipelago ^5,26^ and are critical in the ecology and persistence of some emblematic animal radiations ^3^, *Opuntia* can be considered a keystone element of the Galápagos biota. Understanding the dynamics of the overlooked radiation of prickly pear cacti is therefore essential, not only for revealing the biological impacts of its evolution in other lineages but also for designing conservation strategies to protect the biodiversity of the Galápagos islands as a whole.

**Figure 1.**
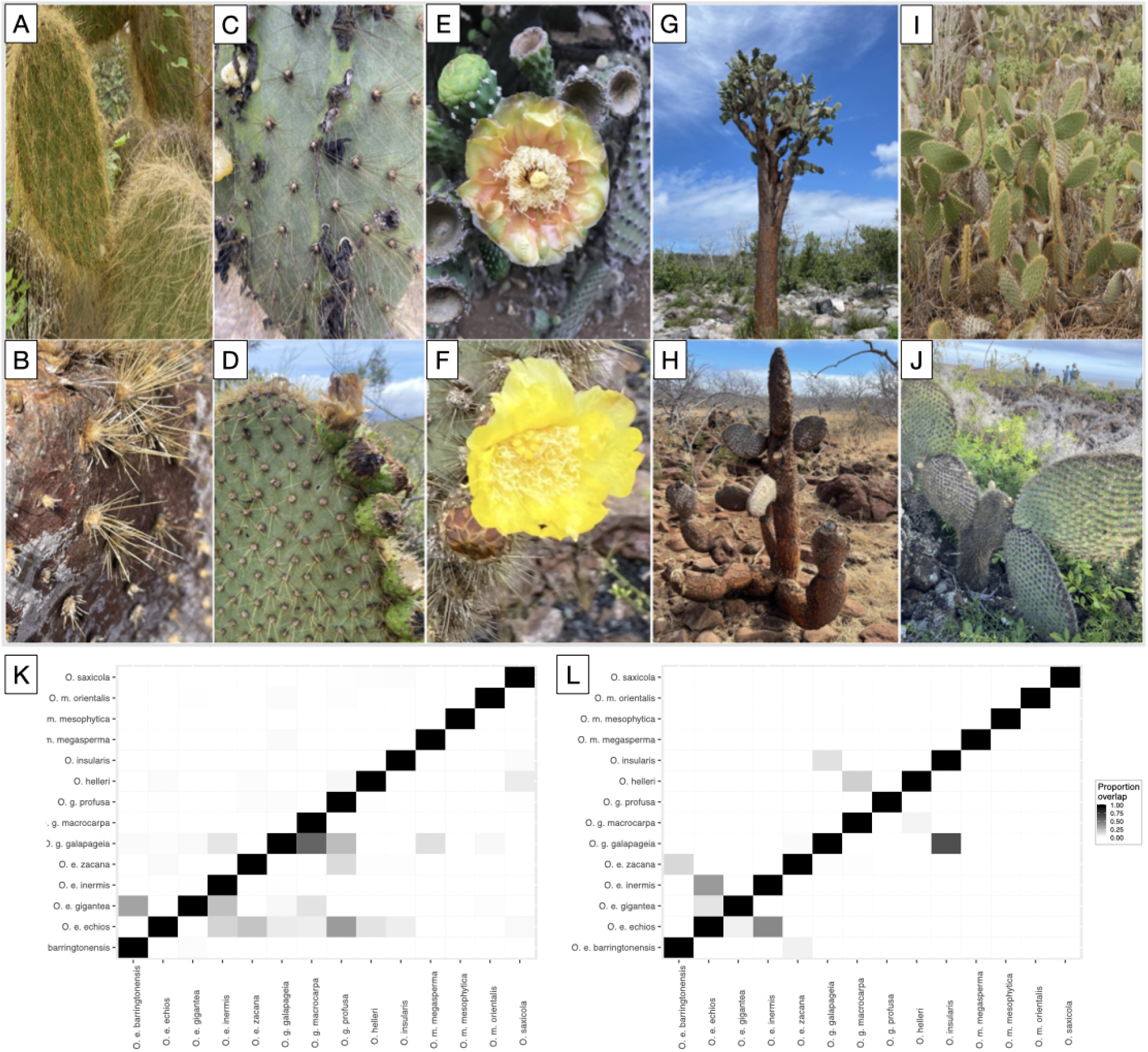
*Opuntia* vary extensively in phenotypic and ecological traits across the Galápagos islands. Spines: (A) *Opuntia megasperma var. megasperma* from Floreana (photo J.A.C.), (B) *O. echio*s *var. zacana* from Seymour (photo G.R.T), (C) *O. galapageia var. galapageia* from Santiago (photo G.R.T), (D) *O. saxicola* from Isabela, Volcan Chico (photo J.E.M); Flowers: (E) *O. megasperma var. mesophytica* from San Cristóbal (photo G.R.T), (F) *O. saxicola* from Isabela, Volcan chico (photo J.A.C.); Habit: (G) *O. echios var. barringtonensis* from Santa Fe (photo J.E.M), (H). *O. echios var. zacana* from North Seymour (photo G.R.T), (I) *O. helleri* from Wolf (photo J.A.C.), (J) *O. insularis* from Isabela (photo G.R.T). (K) Pairwise overlap among hypervolumes describing *Opuntia* taxa in the Galápagos in multidimensional vegetative morphospace (8 traits). Most of the overlap occurs between subspecies of the same species. (L) Pairwise overlap among hypervolumes describing *Opuntia* taxa in the Galápagos in multidimensional reproductive morphospace (5 traits). Most of the overlap occurs between subspecies of the same species

### Genomic lineages of *Opuntia* in the Galápagos

The prickly pear cacti in the Galápagos islands display outstanding eco-phenotypic variation, including 4-fold differences in plant height, 100-fold differences in seed size and seed number per fruit, and 5-fold differences in seed hardness ^24,27^. In addition, species and populations vary extensively in the size of pads as well as the density and morphology of spines ^5^ (Fig. 1). This pronounced phenotypic variation has led to controversies around the nature of species in Galápagos *Opuntia*. Six species are recognized, with three species further subdivided into multiple varieties, for a total of fourteen taxa ^21^. Notably, each recognized taxon is virtually restricted to a single large to mid-size island, with the exception of *O. helleri* which is widespread across multiple islands. Previous work based on the study of variation in eight allozymes, ten microsatellites, and two sequenced loci suggested that despite the remarkable phenotypic and ecological variation of *Opuntia* in the Galápagos, there is exceptionally low genetic variation across this radiation ^20–22^. Therefore, whether each of the different recognized taxa represents independently evolving lineages remains controversial.

To examine the population genetic structure of *Opuntia* in the Galápagos islands, we used 3RAD sequencing ^28^ on 37 individuals representing seven lineages and eight islands, and 2 individuals belonging to an outgroup lineage. Overall, we found that while there is some genetic differentiation among cacti populations across the archipelago, such differentiation is not highly pronounced. The first two principal-component axes ^29^ accounted for only 19.5% (PC1 10% of variance explained; PC2 9.5% of variance explained; Fig 2A) of the variation in the multivariate genomic space across the Galápagos islands. Using up to four principal-component axes accounted for 32.5% of the variation (Fig 2A). Together, these findings indicate that the genomic differentiation of cacti populations across the Galápagos islands, while low, nonetheless reveals some geographic patterns. For instance, PC1 and PC2 broadly separated populations along latitude (Floreana and San Cristóbal separate from Pinta and Genovesa, and Santa Cruz separate from Wolf) while populations at the center of the archipelago (Santiago and Isabela) clustered together in PC space (Fig 2A). Genomic ancestry analyses ^30^ also revealed low genetic differentiation of populations across the archipelago (Fig. 2B). The optimal partitioning of the data indicated the existence of two demes with all the individuals displaying mixed ancestry, irrespective of island of origin or taxonomic assignment (Supplementary Figure S1). This suggests considerable admixture or recent ancestry for all sampled populations.

**Figure 2.**
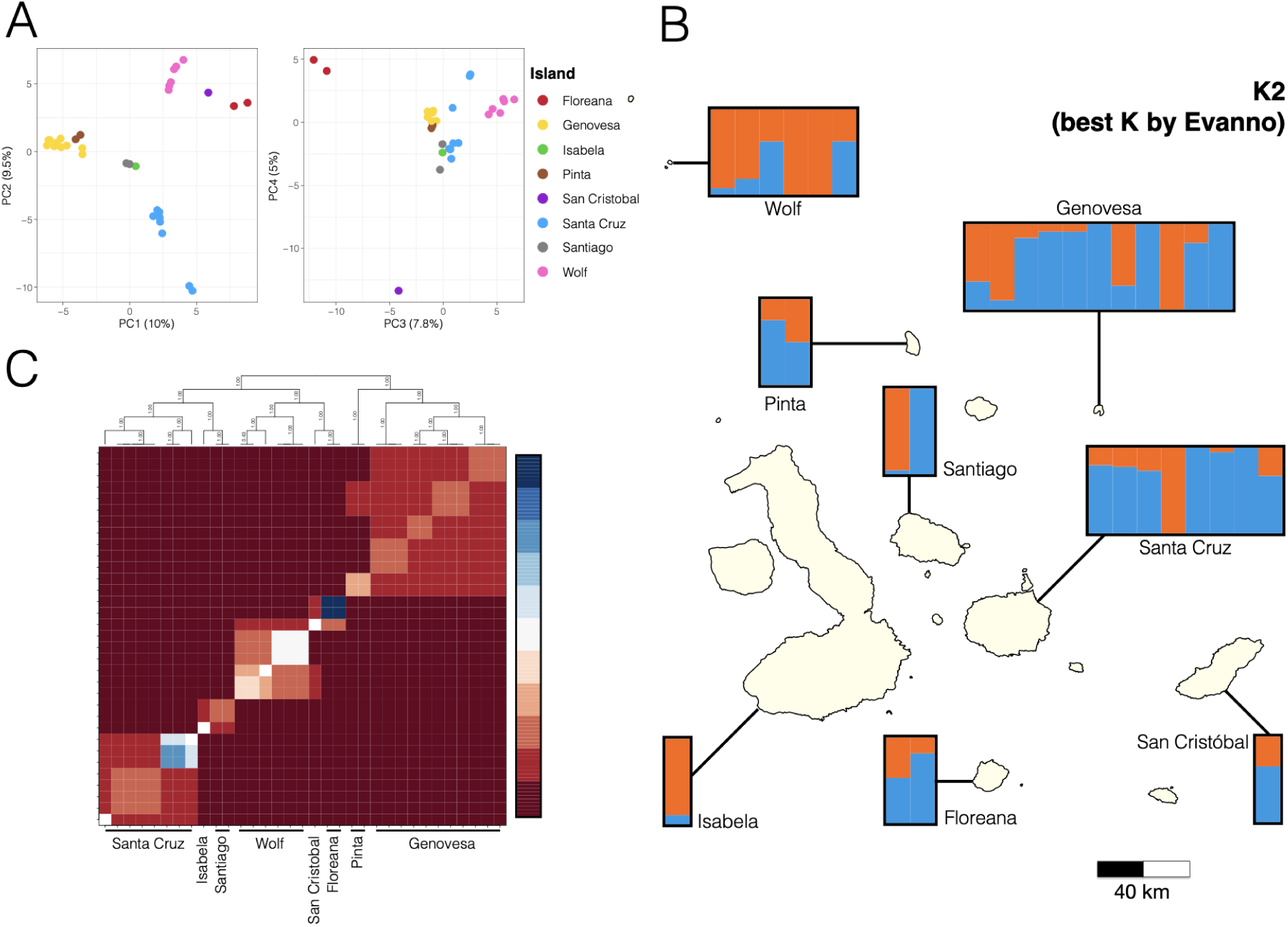
Despite extensive ecophenotypic variation, *Opuntia* shows relatively low genetic differentiation across the Galápagos islands, yet there is a marked geographic structure. (A) Principal component analysis (PCA) to reduce multidimensional genomic space into fewer dimensions. The left panel shows the space defined by PC1 (10% of the variance explained) and PC2 (9.5% of the variance explained), the right panel shows the space defined by PC3 (7.8% of the variance explained) and PC4 (5% of the variance explained). (B) Genomic ancestry shows the optimal partition of the data into two genetic demes shown across islands. (C) Fine-scale population structure based on nearest-neighbor relationships across all individuals sorted by island. Five clusters are readily detected corresponding to single islands or island pairs in close geographic proximity. Further substructure exists within some clusters. Cold colors (blue, top of color legend) correspond to closely related individuals, while warm colors (red, bottom of color legend) correspond to more distantly related individuals.

To characterize fine-scale population structure, we harnessed the power and resolution afforded by RAD-seq data to infer recent shared ancestry at multiple successive SNPs (haplotypes) based on nearest-neighbor relationships ^31,32^. Clustering results readily revealed the presence of five populations, with marked substructure suggested in some of the populations (Fig 2C). The five populations corresponded to groups of samples from single islands or island pairs in close geographic proximity, namely Santa Cruz, Isabela + Santiago, Wolf, Floreana + San Cristobal, and Pinta + Genovesa, with further substructure within island pairs matching single islands (e.g., Isabela and Santiago, Floreana, and San Cristobal). Additionally, our results showed heterogeneous degrees of differentiation of samples between and within all populations, with some localities highly differentiated from samples at other localities (e.g., all samples from Floreana and a subgroup of samples within Santa Cruz). Notably, the highly differentiated subpopulations in Santa Cruz corresponded to the two varieties of *O. echios* present on the island. Overall, these results suggest shallow genetic divergence across populations with limited gene flow and highly restricted geographically.

In sum, our findings show that while genetic differentiation of prickly pear cacti in the Galápagos archipelago is low, there is a marked geographic pattern of genetic structure by islands (Fig 2A, C). This is consistent with the current taxonomic hypothesis that almost every major island harbors a unique taxon, a hypothesis worthy of further investigation with increased genomic sampling in combination with explicit analysis of phenotypic variation to examine in detail the nature of *Opuntia* species in the Galápagos ^33,34^. Conversely, it is plausible that the geographic pattern of genetic variation displayed by *Opuntia* reflects mostly intraspecific variation and the early stages of a radiation in prickly pear cacti across the Galápagos archipelago, likely fueled by phenotypic plasticity and divergent selection resulting from island-specific environments and biotic interactions. Indeed, it has been proposed that environmental variation is an important driver of phenotypic expression in cacti more broadly ^35^ and that biotic interactions may contribute specifically to explain patterns of phenotypic diversity across *Opuntia* species in the Galápagos ^24,27^. In the future, testing this intriguing hypothesis with increasing rigor would shed light on the genomic underpinnings of phenotypic plasticity and organismal adaptation ^36,37^.

### Radiation across the Galápagos archipelago

The prickly pear cacti are a prominent component of the Galápagos flora. They are widespread and occur on all major islands (> 10 km^2^) and most islets. Although there is considerable doubt about the most closely related species on the mainland ^38^, it is likely that *Opuntia* colonized the Galápagos only once drafting from the nearby continent via ocean currents as most other native plants of the archipelago ^39^. Once in the archipelago, *Opuntia* diversified across islands in multiple environments, ranging from coastal dry open scrublands to higher elevation wet forests with closed canopies ^26^. Yet, the colonization history and the biological mechanisms enabling inter-island dispersal are poorly understood.

To reconstruct the radiation of *Opuntia* throughout the Galápagos, we grouped individuals by island and used islands as units of analysis. This approach is consistent with our results on genetic structure (see above) and current taxonomy, as well as a reasonable approach to studying the colonization of organisms across archipelagos. Owing to the dynamic paleogeography of the Galápagos, including, but not limited to, volcanism linked to plate tectonics, the existence of ancient proto-islands, the subsidence and growth of extinct and extant islands, and changes in the connectivity and isolation of the current islands ^40^, combined with the vagility of the seeds dispersers of cacti plants ^5,41^, nearly any of the islands could be the source of colonization for other islands, and the sequence of dispersal routes could be extraordinarily complicated. Nonetheless, our results revealed two notable patterns.

First, phylogenetic reconstructions suggest a recent radiation of prickly pear cacti in the Galápagos, as seen in the tangled phylogenetic network (Fig. 3A), the short internal branches in the species tree (Fig. 3B), and the topological tree discordance using different subsets of loci (Fig. 3C). Additionally, we discovered a remarkably low number of private variants per island in our dataset (Table 1). Together, these findings are consistent with a history of diversification influenced in part by the recent dynamic geological history of the archipelago ^42^. Although the estimated maximum age of the oldest present Galápagos island is around 4 my (San Cristóbal), the bulk of the present islands have a maximum age of about 2.5 my ^21,40^. This age corresponds to the beginning of the Pleistocene, a geological epoch characterized by repeated glacial-interglacial cycles that have modified dramatically the geography and connectivity of oceanic archipelagos around the globe ^43^, including the Galápagos islands. Just within the last 3 My, the time frame of the emergent history of the present Galápagos islands, there have been approximately 36 glacial advances separated by interglacials ^44^. Although the exact imprint of these geo-climatic events is difficult to reconstruct, Geist et al. ^40^ suggest a complex paleogeographic model for the Galápagos archipelago. They propose that only two of the nine islands present 3 Mya still exist today (San Cristóbal and Española). At 2 Mya, Santa Cruz and Floreana were a single large island and 10 other islands existed, many of which are not present today. At 1 Mya, it is likely that a massive central island was an amalgamation of multiple present islands (Santiago, Pinzón, Rábida, Santa Cruz, and Floreana), and several other islands emergent back then are submerged today. Even in more recent times, during the Last Glacial Maximum (approximately 20 Ky), there were many more islands and islets present and there were land bridges connecting three of the present big central islands (Isabela, Fernandina, and Santa Cruz). In sum, such drastic paleogeographic changes clearly affected the connectivity of populations across islands, thus possibly shaping directly the recent radiation of prickly pear cacti in the archipelago ^45^.

**Figure 3.**
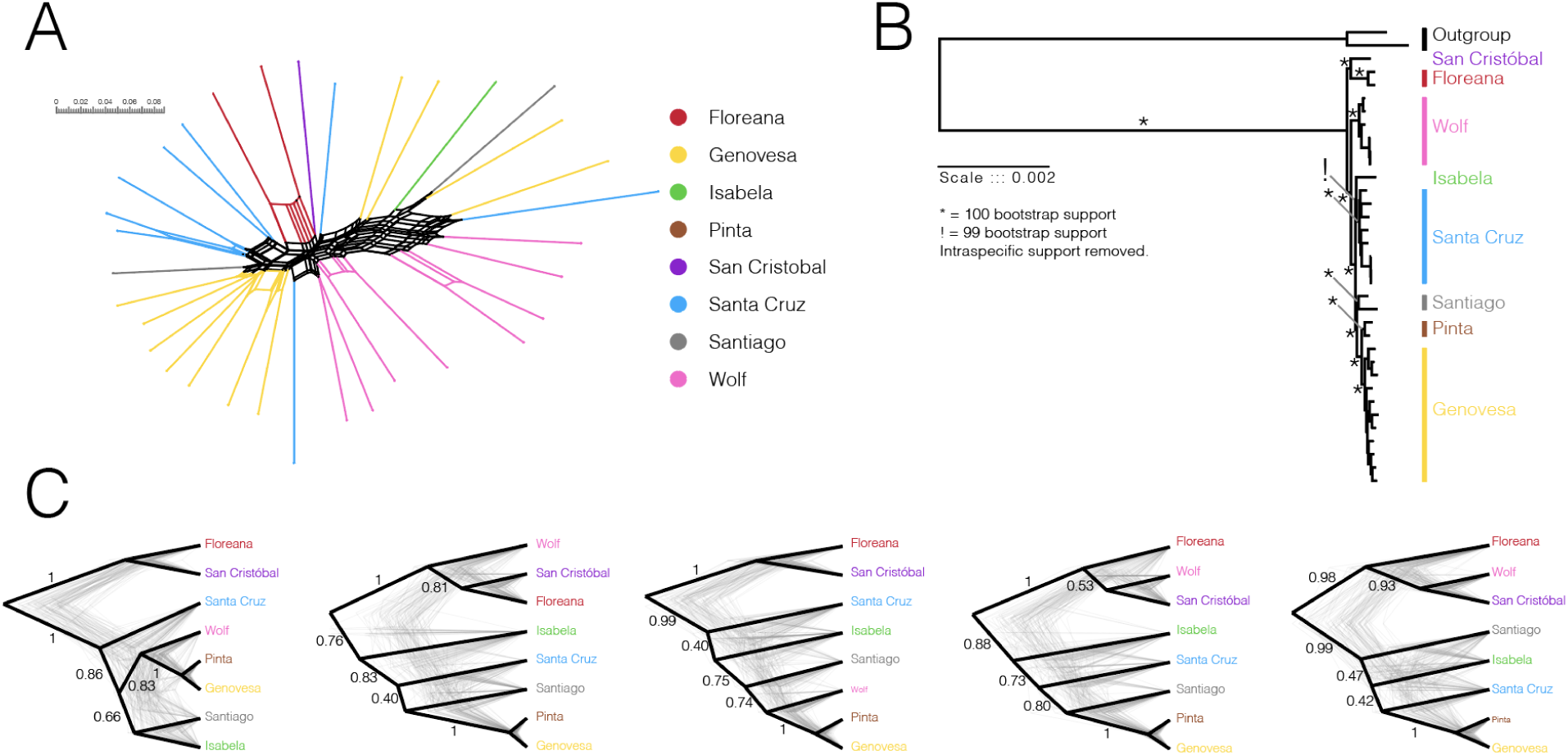
The radiation of *Opuntia* across the Gálapagos archipelago is a recent phenomenon. (A) The phylogenetic network shows tangled relationships among all island populations. (B) The phylogenetic tree among all island populations using a concatenated matrix of all loci shows very short internal branches subtending phylogenetic relationships. (C) The phylogenetic trees among all island populations using a coalescent approach (without concatenation) vary according to the loci included in the analysis. Each panel shows trees inferred with 5,000 random sampled loci. The clouds of trees in the background represent trees sampled from the posterior distribution, the black like represents the Maximum Credibility Tree.

**Table 1.** Private variant analysis shows low differentiation of *Opuntia* across the Galápags. For each island, we report the number of island-specific variants. Private variants correspond to alleles that only exist on a particular island. This analysis is based on a dataset of 52,869 total SNPs.

| Island | Private Variants | % Private Variants |
| --- | --- | --- |
| Floreana | 1,805 | 3.4% |
| Genovesa | 2,831 | 5.4% |
| Isabela | 891 | 1.7% |
| Pinta | 738 | 1.4% |
| San Cristóbal | 949 | 1.8% |
| Santa Cruz | 3,916 | 7.4% |
| Santiago | 1,314 | 2.5% |
| Wolf | 1,661 | 3.1% |
| **Total** | **14,105** | **26.7%** |

Alternatively, our phylogenetic results and the distribution of private variants could be the outcome of gene flow between geographically isolated populations, perhaps facilitated by pollen and seed dispersers ^24^ and weak isolating barriers. For instance, we found that the population from San Cristóbal is sister to the population from either Floreana or Wolf, depending on the loci used to infer phylogenetic relationships (Fig. 3B, C). Such a pattern could result from differential gene flow among populations that are separated by more than 300 kilometers like in the case of San Cristobal and Wolf.

To test formally for genetic admixture between populations in our data set, we used the F4 statistic ^46^. We detected excess allelic sharing between the populations from San Cristóbal and Wolf, Genovesa and Santa Cruz, and Wolf and the grouped populations from Genovesa and Pinta, indicating ancient admixture between all these populations (Fig. 4). Previous studies have suggested that as mainland *Opuntia* species have weak reproductive barriers ^21^ and because the two sympatric taxa of *Opuntia* present in Santa Cruz island show little to no signal of genetic differentiation based on microsatellite data ^22,45^, gene flow between lineages might be widespread across the Galápagos *Opuntia*. Our results are in general agreement with this idea, yet our evidence for admixture shows that gene flow is more prevalent between certain lineages than others. It remains unclear whether the instances of admixture we report here correspond to cases of hybridization or introgression and it will require more detailed analyses of patterns of reproductive isolation and ecological differentiation between lineages.

**Figure 4.**
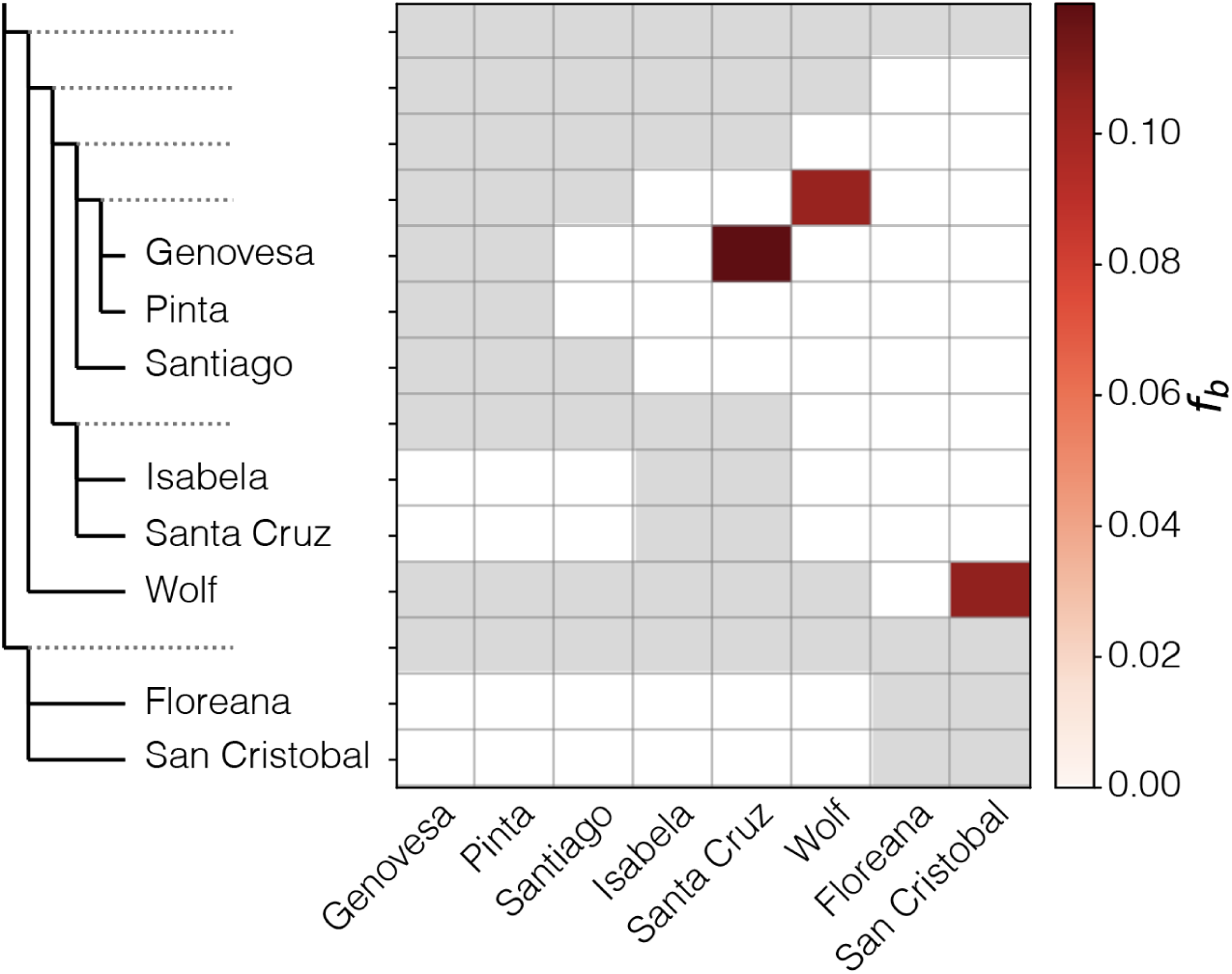
Genetic admixture between populations of *Opuntia* across the Galápagos. The phylogenetic tree obtained using the concatenated matrix (see Figure 3B) is shown on the left in an “expanded” form, so that each branch, including internal branches, points to a corresponding row in the matrix with the inferred *f*-branch statistics. The values in the matrix refer to excess allele sharing between the branch identified on the expanded tree on the y-axis (relative to its sister branch) and the population on the x-axis. Warm colors (red, top of the color legend) refer to excess allele sharing.

Likewise, whether genetic admixture fuels or constraints differentiation is not known. Investigating the extent to which gene flow could increase the amount of genetic variation in populations to promote adaptation to local conditions will require identification of the loci underlying the traits of interest and characterization of their evolutionary histories ^47^.

The second notable phylogenetic result concerns the geographic structure of the radiation of Galápagos *Opuntia*. Both the concatenated and coalescent-based species tree analyses recovered consistently the oldest divergence of the radiation to be the separation between the populations from San Cristóbal and Floreana, and sometimes Wolf, from the remaining populations (Fig. 3B, C).

Because San Cristóbal is the oldest current island (2.4 - 4.0 my) ^40^, the deep divergence event separating the *Opuntia* population from San Cristóbal is consistent with the history of the islands. However, the age of Floreana (1.5 - 2.3 my) is on par with the ages of several other current islands included in our dataset, such as Santa Cruz (1.1 - 2.3 my) and Wolf (1.6 - 1.7), as well as with the ages of Pinzón (1.3 - 1.7 my), Rábida (1.3 - 1.6 my), and Santa Fé (2.9 -2.9) not included in our sampling. Furthermore, Floreana was part of a larger landmass that included Santa Cruz at least 2 mya ^40^. Therefore, although the deep split separating the *Opuntia* population from San Cristóbal and the other islands is consistent with the age of this island, it is unclear if the colonization of Floreana involved a stepping stone dispersal via other older islands such as Española, Santa Fé, or Pinzón, for which we are missing *Opuntia* samples. Including samples from such islands in combination with explicit biogeographic analyses is required to shed light on this possibility.

Sister to the San Cristóbal and Floreana (and sometimes Wolf) clade, the relationships among the remaining populations are broadly consistent with the estimates of island ages and geographic proximity (Fig. 3B, C). The populations from Genovesa and Pinta are sister to each other and are nested within a clade which includes the populations from Santiago, Santa Cruz, and Isabela.

Genovesa and Pinta are some of the youngest islands in our dataset, with ages ranging between 0.3 - 0.8 my and 0.7 - 0.8 my, respectively ^40^. Because these islands are also far apart (∼80 km) from the centrally located islands (e.g., Santa Cruz, Isabela, Santiago) and have never been connected to other islands in the archipelago in geological time, our result suggests a recent, long-distance dispersal colonization of *Opuntia* to the islands of Genovesa, Pinta, and possibly Marchena for which we are missing samples in the current study. The ages of Santa Cruz and Santiago islands range between 1.1 -2.3 my and 0.8 - 1.4 my, respectively. In contrast to other current islands included in this study, these two islands have been part of the same landmass at some point over the last million years ^40^. Isabela is one of the youngest islands of the archipelago, with an age estimate of 0.5 - 0.8 my. Further, Isabela is in extremely close geographic proximity to Santa Cruz and Santiago, and it is likely that there was a landbridge connecting Isabela and Santa Cruz during the Last Glacial Maximum (20 ky) ^40^. Therefore, our results are consistent with the colonization of the centrally located islands (Santigo, Santa Cruz, and Isabela) from an older island (extinct or extant), with subsequent, repeated movements of plant propagules among all the central islands, likely facilitated by the shared geology and close geographic proximity. Evidence for this scenario comes from the uncertain and incongruent relationships among the populations of these islands (Fig 3C).

Intriguingly, the relationship between the *Opuntia* population from Wolf and populations from all other islands is the most uncertain. Although such uncertainty is largely explained by instances of gene flow (Fig. 4), Wolf is a small (1.3 km^2^) and distant island (>150 km apart from other islands with the exception of Darwin island) in the northwest corner of the archipelago, and it is not known how *Opuntia* could have colonized this island. Wolf is approximately as old as Floreana (1.6 - 1.7 my; ^40^), hence it might have been available for colonization for a long period of time, yet how *Opuntia* propagules reached this remote location in the first place remains elusive. While the direction of sea currents in the Galapagos archipelago is westward and northwestward ^23^ and this could have facilitated the dispersal of *Opuntia* to Wolf, it is more likely that animals helped disperse these plants. The presence in Wolf Island of birds known to disperse *Opuntia* seeds elsewhere, such as Darwin’s finches (*G. septentrionalis* and *G. magnirostris*: JAC pers. comm.) and mockingbirds, could support this colonization hypothesis. Lava lizards on other islands also act as seed dispersers for *Opuntia* (*O.echios* in Santa Cruz and *O. galapageia* in Pinta) ^48^, but no lava lizards are currently found in Wolf or Darwin islands. A general trend of unidirectional gene flow and historical migration on several species within the archipelago ^12,49–52^ resulting from prevailing south–south-east trade winds in the Galápagos ^53–55^ could explain the colonization of *Opuntia* propagules or seeds dispersed by animals on these northern islands. Complementing our current sampling with populations from all other current Galápagos islands will certainly help clarify the biogeography and potential dispersal routes of *Opuntia* across the archipelago.

### A potential genomic link to adaptive phenotypes

One of the most remarkable attributes of the Galápagos prickly pear cacti radiation is the extraordinary phenotypic diversity they display in vegetative and reproductive traits (Fig. 1). Previous studies have suggested that both environmental and biotic factors may be responsible for such striking phenotypic diversity. In examining the patterns of morphological variation of *Opuntia* in three centrally located islands (Santa Fé, Santa Cruz, and Pinzón), Racine and Downhower ^41^ suggested that variation in tree height-diameter growth and pad production was largely explained by the combined effects of inter and intraspecific competition for light and wind velocity. In addition to the extensive variation observed in vegetative growth form, these authors also proposed that the pronounced variation in seed size and seed coat observed among island populations was related to seed dispersal and predation. While mockingbirds, tortoises, and iguanas disperse seeds by consuming either the fruits or the fleshy aril around the seed, the cactus finch (*Geospiza scandens*) is a major seed predator that cracks the seed coat to extract the endosperm ^27^. Hence, it is likely that seed predation is an important factor in the diversification of *Opuntia*, and the cactus finch is a prime agent of selection. Subsequent studies are consistent with this hypothesis and show that the interaction between Darwin’s finches and prickly pear cacti extends, in fact, to multiple species of ground finches and cacti ^24^. The intricate relationship between these two lineages is largely modulated by environmental changes such as increased and prolonged periods of drought, which affect plant survival and thus seed availability. Long-term studies of Darwin’s finches populations have shown how such changes have had drastic effects on the morphology of the beaks of the finches, which are critical for seed manipulation ^3^. Thus to some extent, it is plausible there is some degree of coevolution between cactus and *Geospiza* finches in the Galápagos mediated by flowers and seed features which requires rigorous study ^2,18,19,56–59^

In order to detect potential genetic contributions linked to phenotypic variation in *Opuntia* across islands, we conducted parallel scans of genetic differentiation (fixation index, F_ST_) between pairs of populations with more than five individuals, namely Genovesa, Santa Cruz, and Wolf. We selected the 5% divergent RADseq tags and mapped these to *Arabidopsis thaliana* gene sequences using conservative cut-offs (Supplementary Table S2). Three genomic regions stood out, as these showed differentiation across all pairwise comparisons. The first region was the *CLocus_1489*, which is associated with the MATE efflux protein family (TT12 or ATTT12). This protein family encodes a multidrug and toxic compound extrusion (MATE) transporter involved in seed coat pigmentation ^60^ and transportation of proanthocyanidins which are pigmentation proteins ^61,62^. The second region was the *CLocus_59059*, which is likely involved in DNA replication initiation and elongation ^63^. The third region was detected as two loci, *CLocus_4279* and *CLocus_4683*, which are associated with a periderm regulator (RIK) ^64^. The periderm is a critical tissue protecting the vascular tissue from biotic and abiotic stresses. When considering each pairwise comparison separately, we detected differentiation in genomic regions associated with environmental stress, including temperature, drought, and salt, genomic regions associated with light responses and circadian clock, as well as genomic regions associated with plant pigmentation, among others (Supplementary Table S2).

Although significant differentiation detected via F_ST_ scans does not necessarily imply differentiation via natural selection ^65^ and further functional validation is desirable ^66^, some results are noteworthy. Genomic differentiation linked to chemical compounds involved in seed pigmentation might be related to seed predation, supporting the hypothesis of eco-evolutionary feedback between *Opuntia* and Darwin’s finches ^24^. Alternatively, it is plausible that variable pigmentation could confer better camouflage to different soil types, a scenario congruent with the proposal of seed dormancy in *Opuntia* ^67^. Further sampling and experimental data are necessary before these hypotheses can be confronted confidently. Genomic differentiation associated with regulation and development of the periderm as well as with multiple regulators involved in environmental stresses might be related to the harsh conditions that characterize the Galápagos islands, and possibly biotic interactions. The Galápagos islands lie directly on the equator, where sun exposure may be a major environmental stress for island dwellers. One of the Darwin daisies from the Galápagos, *Scalesia atractyloides*, shows signatures of selection along the genome associated with light reception and growth regulation ^7^.

Additionally, some *Opuntia* plants grow on harsh volcanic soils and some individuals are exposed constantly to salt spray, indicating that prickly pear cacti deal with considerable sustained abiotic stresses. Likewise, selective pressures due to biotic stress may occur from tortoises, which are known to be the main herbivores of prickly pear cacti in the Galápagos ^5^. Taken together, our results suggest that a combination of environmental and biotic selective pressures may have left an imprint on the cacti’s genomes, an observation worthy of closer investigation to unravel the genomic underpinnings and molecular mechanisms of the *Opuntia* radiation in the Galápagos archipelago.

### Conclusions

This study presents genomic evidence of decoupling between phenotypic and genomic differentiation in the Galápagos *Opuntia*. While cactus populations from different islands show extreme variation in vegetative and reproductive traits (Fig. 1), the evidence for concomitant genomic variation is weak (Figs. 2, 3). This is consistent with the hypothesis that these cacti are at the early stages of a radiation and that gene flow between populations may facilitate local adaptation ^68^. Alternatively, it is plausible that phenotypic plasticity precedes genetic change and is the source of adaptive divergence ^69^. Further geographic sampling, deeper genome sequencing, and experimental data are critically needed before these hypotheses can be confronted confidently and the biological mechanisms underpinning this radiation are better understood. The observed patterns of parallel differentiation in genomic regions associated with seed pigmentation and periderm development suggest seed predation and abiotic stresses as potential mechanisms of adaptation that may be targets of selection between populations. Further study is needed to determine whether these patterns are widespread across all islands in the archipelago to understand how these genomic changes contribute to plant fitness and how selection acts during local adaptation.

## MATERIALS AND METHODS

### Taxon Sampling and DNA Extraction

Samples were collected from 8 islands across the Galapagos archipelago (Figure 2; Supplementary Table S1). The localities selected represented most of the islands across the geographic range of *Opuntia* in the Galapagos. We assigned all specimens to species following the taxonomy of Helsen et al. ^21^ (Supplementary Table S1). We collected approximately 1-2 cm^2^ of pad tissue for each specimen, stored it in silica gel, and then transported it to the University of California, Los Angeles for further analyses. Genomic DNA extraction and purification were performed following a modified version of the CTAB extraction protocol ^70,71^ that incorporates a pre-wash step [@li2007optimized] to aid in the removal of polyphenols and proteins prior to extraction. Genomic DNA quantification was performed using a Qubit fluorometer v.3.0 (Invitrogen by Thermo Fisher Scientific, Carlsbad, CA, USA) and an Agilent 2200 TapeStation (Agilent Technologies, Santa Clara, CA, USA).

### Library Preparation and Sequencing

To generate sequence data, we prepared quadruple-indexed, triple-enzyme RADseq libraries using the *EcoRI*, *XbaI*, and *NheI* restriction enzymes ^28^. Prior to sequencing, size selection for fragment length of 375-525 bp was performed on a PippenPrep (Sage Science, Beverly, MA, USA). Libraries were pooled and sequenced in one lane of 100PE sequencing on the Illumina HiSeq4000 Sequencing Platform at the Broad Stem Cell Research Center at the University of California, Los Angeles.

### Bioinformatics and SNP discovery

We demultiplexed, removed adapters, and filtered reads for data quality using the iPyrad package v 0.9.72 ^72^. Because of the potential polyploid nature of the Galápagos *Opuntia* ^38^, we carefully explored and processed the data. Specifically, we ran Stacks v2.60 ^73^ using different M/N combinations as suggested by ^74^ to optimize rad-assembly parameters. This involved running stacks 10 times using -M (number of mismatches allowed between stacks within individuals) -N (number of mismatches allowed between stacks between individuals) values between 1 and 10. We ran this protocol four times, specifying different population maps (i.e. groups of populations), namely by islands, by species, by islands excluding outgroups, and by islands excluding outgroups. This led to an optimal combination of -M / -N of 2 for analysis involving the outgroup and -M 1 -N 1 for analysis not including the outgroup.

Because samples with a high degree of missing data tend to compromise the quality of the final RADseq variant dataset, we followed the protocol established by ^75^ to remove “bad apples”. This led to the removal of three outliers with high levels of missing data, resulting in a final dataset of 33 individuals of Galápagos *Opuntia* and two outgroup specimens. For the final dataset, we ran stacks with and without outgroups, specifying -R 0.8 (minimum percentage of individuals across populations required to process a locus).

### Population genetics and phylogeographic analyses

We performed a phylogenetic reconstruction and a Dsuite analysis using the Stacks run with outgroups. For the phylogenetic reconstruction, we retrieved a collection of FASTA rad loci (--fasta-samples). Since we had a dataset of 35 individuals (70 alleles), we kept loci with more than 59 data points (30 individuals), retaining a total of 10,484 loci (2,134,262 bp). We obtained a consensus sequence between the two loci using EMBOSS v6.6.0 consambig as done in ^76^ and concatenated the consensus loci using FASconCAT-G v1.04 (https://github.com/PatrickKueck/FASconCAT-G/). We used the concatenated loci to run IQ-TREE v2.1.3 ^77,78^ specifying 1,000 ultrafast bootstraps.

We ran Principal Component Analysis (PCA), STRUCTURE, a private allele analysis, fineRAD, Splitstree, and SNAPP with the dataset without outgroups. For the PCA and STRUCTURE analysis, we obtained a vcf file where linkage was reduced by specifying only a single and random SNP per radtag (--write-random-snp), and then cleaned the vcf for coverage and for missing data as specified for the F4 analysis, obtaining a total of 23,566 SNPs. To run the PCA, we used the libraries vcfR ^79^, adegenet ^80^, and ggplot2 ^81^. For STRUCTURE analysis we used plink ^82^ to convert the VCF into a structure file, and ran STRUCTURE v2.3.4 ^30^ specifying Ks between 2 and 10 (Burn-in of 100,000 and an MCMC chain of 100,000). For plotting the results of this analysis, we used the online server CLUMPAK ^83^ and its plug-in which calculates the Evanno method to determine the optimal K ^84^. For the remaining analyses (private allele analysis, SplitsTree, and SNAPP) we used a VCF where filtering included only coverage (--maxDP 200 --max-meanDP 200 --minDP 10 --min-meanDP 10), missing data (--max-missing 0.25) and loci shared (-R 0.8). For the private alleles, we used vcfR and the function ‘genetic_diff’ to calculate measures of genetic differentiation (method = nei) by population, retrieving a table with a description of heterozygotes and homozygotes, which was then parsed to determine alleles private to the different populations. We then ran fineRADstructure v0.3.2 ^32^ using default settings and annex plotting scripts (https://www.milan-malinsky.org/fineradstructure). For the SplitsTree analysis, we converted the vcf into a nexus file using the script vcf2phyl.py (https://github.com/edgardomortiz/vcf2phylip), and used SplitsTree v5 to estimate a network ^85–87^.

For SNAPP, we subsampled the VCF to 4,000 random SNPs and ran SNAPP as included in BEAST2^88,89^. For the F4 analyses, we used Dsuite ^90^. This involved retrieving a vcf from Stacks (-R 0.8), and filtering it further by coverage (--maxDP 200 --max-meanDP 200 --minDP 10 --min-meanDP 10) and missing data (--max-missing 0.25) using vcftools ^91^. We then ran the algorithm Fbranch of Dsuite specifying the tree obtained above and plotting it using a collection of ruby scripts from Michael Matschiner (https://github.com/mmatschiner/tutorials/blob/master/analysis_of_introgression_with_snp_data/ <u>README.md</u>).

### Functional genetics

For the functional analysis, we ran FST with a window size of 1,000 bp and window step of 1,000 bp (independent for each rad locus) using the Weir and Cockheram estimate as implemented in VCF tools ^91^. Using the weighted estimate, we extracted the top 5% most differentiated between Genovesa vs Wolf (446 loci at the 5% cut-off), Genovesa vs Santa Cruz (490 loci at the 5% cut-off), and Santa Cruz vs Wolf (466 loci at the 5% cut-off), and used blast 2.9.0 to compare with the *Arabidopsis thaliana* protein list (https://www.arabidopsis.org/download_files/Proteins/TAIR10_protein_lists/TAIR10_pep_201012 <u>14</u>). To do this, we created a protein database (makeblastdb -dbtype prot) and ran blastx with the top 5% outlier radtags. We then extracted e-values below 0.001 and kept only amino acids with a sequence length of 20. We searched the literature for functional information associated with each locus hit.

### Phenotypic variation

We used the most recent taxonomic monograph of Galpágos *Opuntia* to tabulate the minimum and maximum values reported for 13 phenotypic traits used to describe and delimit each of the 14 recognized taxa ^26^. Vegetative traits included plant height, pad length, pad width, pad thickness, areoles width, distance between adjacent areoles, number of spines per areole, and spine length.

Reproductive traits included fruit length, fruit width, seed length, seed width, and seed thickness. The combination of minimum and maximum values of these traits delimits a hypervolume of *n* dimensions (*n* = the number of phenotypic traits) in the phenotypic space corresponding to each species. We constructed separate hypervolumes for vegetative traits (n = 8) and reproductive traits (n = 5) because some *Opuntia* species seem to differ more in one kind of trait than the other. To determine the distinctiveness of each taxon, we log-transformed all data and estimated the pairwise asymmetric proportion of overlap of all vegetative and reproduction hypervolumes. We used the library *geometry* ^92^ in R to carry out this analysis.

### Data and Code Availability

All sequence data have been deposited to the Sequence Read Archive (NCBI): BioProject ID PRJNA1022515 (https://www.ncbi.nlm.nih.gov/bioproject/1022515). All processed data files to generate results are available at https://github.com/zapata-lab/ms_opuntia_genomics All code to run the analyses is available at: https://github.com/zapata-lab/ms_opuntia_genomics

## SUPPLEMENTAL INFORMATION

Supplemental Information includes:

- Summary of the study in Spanish.
- Supplementary Figure S1: alternative results to partition the data into demes (K=2, K=6, K= 8).
- Supplementary Table S1: taxon sampling metadata
- Supplementary Table S2: results of scans of genetic differentiation (fixation index, F_ST_) between pairs of populations with more than five individuals.

## ACKNOWLEDGMENTS

The authors thank the Galápagos National Park. All plant materials collected and used in this investigation were regulated by permits no. MAE-DNB-CM-2016-0041-M-0002 and PC 87-22 provided by the Galápagos National Park-Ministerio del Ambiente, Ecuador to G.R.-T and J.C. This work was partially funded by a UCLA - Faculty Research Grant to F.Z. This work used computational and storage services associated with the Hoffman2 Shared Cluster provided by UCLA Institute for Digital Research and Education’s Research Technology Group, and the Norwegian national infrastructure for high-performance computing and storage. J.C. thanks Angel Rivera-Colon and Nicolas Rochette for invaluable comments on RADseq analyses of polyploids.

## AUTHOR CONTRIBUTIONS

Conceptualization, F.Z., J.C., J.E.M., J.A.C., and G.R.-T.; Investigation, F.Z., J.C., D.M, C.L.H., B.B.T., J.E.M, K.R.R., J.A.C., and G.R.-T.; Resources, F.Z., J.C., J.E.M, J.A.C, and G.R.-T.; Data Curation, F.Z., D.M, C.L.H., J.E.M, and K.R.R.; Writing – Original Draft, F.Z. and J.C.; Writing – Review and Editing, all authors; Visualization, J.C.; Supervision and Project Administration, F.Z. and G.R.-T.; Funding Acquisition, F.Z.

## DECLARATION OF INTERESTS

The authors declare no competing interests.

## Notes

https://www.ncbi.nlm.nih.gov/bioproject/1022515

### Competing Interest Statement

The authors have declared no competing interest.

